# *In vivo* CRISPR screens identify novel virulence genes among proteins of unassigned subcellular localization in *Toxoplasma*

**DOI:** 10.1101/2024.01.28.577556

**Authors:** Yuta Tachibana, Miwa Sasai, Masahiro Yamamoto

**Author notes:** Correspondence: Masahiro Yamamoto.

## Abstract

The research field to identify and characterize virulence genes in *Toxoplasma gondii* has been dramatically advanced by a series of *in vivo* CRISPR screens. Although subcellular localizations of thousands of proteins were predicted by the spatial proteomic method called hyperLOPIT, those of more than 1000 proteins remained unassigned and their essentiality in virulence was also unknown. In this study, we generated two small-scale gRNA libraries targeting approximately 600 hyperLOPIT-unassigned proteins and performed *in vivo* CRISPR screens. As a result, we identified several *in vivo* fitness-conferring genes that were previously unreported. We further characterized two candidates, TgGTPase and TgRimM, which are localized in the cytoplasm and the apicoplast, respectively. Both genes are essential for parasite virulence and widely conserved in the phylum Apicomplexa. Collectively, our current study provides a resource for estimating the *in vivo* essentiality of *Toxoplasma* proteins with previously unknown localizations.

**IMPORTANCE:** *Toxoplasma gondii* is a protozoan parasite that causes severe infection in immunocompromised patients or newborns. *Toxoplasma* possesses more than 8000 genes; however, the genes that determine parasite virulence were not fully identified. The apicomplexan parasites, including *Toxoplasma*, developed unique organelles that do not exist in other model organisms; thus, determining the subcellular location of parasite proteins is important for understanding their functions. Here, we used *in vivo* CRISPR genetic screens that enabled us to investigate hundreds of genes in *Toxoplasma* during mouse infection. We screened approximately 600 parasite proteins with previously unknown subcellular localizations. We identified many novel genes that confer parasite fitness in mice. Among the top hits, we characterized two essential virulence genes, TgGTPase and TgRimM, which are widely conserved in the phylum Apicomplexa. Our findings will contribute to understanding how apicomplexans adapt to the host environment and cause disease.

## INTRODUCTION

An obligate intracellular parasite, *Toxoplasma gondii*, is an important human and animal pathogen that causes life-threatening toxoplasmosis (1). *Toxoplasma* has been the model apicomplexan parasite owing to its easiness of *in vitro* cultivation, genetic tractability, *in vivo* disease model in mice, and various biological assays to investigate the function of the genes (2). As unicellular eukaryotes that adapt to intracellular parasitism, apicomplexans have developed unique secretory organelles (e.g., micronemes, rhoptries, dense granules), which are absent in other well-studied eukaryotic cell systems (3). These secretory organelles are key features of the apicomplexans and discharge various types of proteins that are essential for invasion, replication, and egress (3). Likewise, the apicoplast is a unique non-photosynthetic plastid-like organelle and essential for apicomplexans’ metabolism (fatty acid, heme, iron-sulfur cluster, and isoprenoid synthesis) (4). Therefore, uncovering the subcellular localizations of parasite proteins is crucial to understand the protein functions and parasite pathogenesis mechanism.

Although the parasite protein localization is typically revealed by cell biological methods such as classical immunofluorescence assay (IFA), immuno-electron microscopy, and, recently, ultrastructure expansion microscopy (U-ExM) (5, 6), the parasite spatial proteome has been largely unknown. Recent studies elegantly addressed the issues in *Toxoplasma* (7) and *Cryptosporidium* (8) by a spatial proteomic method called hyperLOPIT (9), yielding plenty of knowledge on parasite subcellular proteome. In a previous study using hyperLOPIT on *Toxoplasma* (7), approximately 3800 proteins were identified in the tachyzoite stage; however, about 1,200 of them are still classified as unassigned subcellular localizations. Moreover, the functions of these genes mostly remained to be elucidated.

In addition to classical forward and reverse genetics, *in vitro* or *in vivo* CRISPR genetic screens have been used to assess the essentiality of the parasite genes in the lytic cycles or virulence (10–15). We previously performed *in vivo* CRISPR screens using small-scaled guide RNA (gRNA) libraries based on hyperLOPIT-assigned localizations (e.g., rhoptries, dense granules, nucleus, endoplasmic reticulum (ER), Golgi apparatus, cytosol, apicoplast, and mitochondrion) and identified novel secreted and non-secreted virulence factors (14). However, the essentiality of hyperLOPIT-unassigned proteins in virulence remains mostly unknown.

In this study, we generated two small-scaled gRNA libraries targeting genes encoding hyperLOPIT-unassigned proteins and performed *in viv*o CRISPR screens in mice. We identified several novel genes that contribute to the parasite’s *in vivo* fitness. Among the top hits, we further focused on two candidates, TgGTPase (TGGT1_277840) and TgRimM (TGGT1_321310). TgGTPase is a guanosine triphosphatase (GTPase) that localizes in the parasite cytosol, and TgRimM is a novel apicoplast-resident protein possessing a ribosome maturation factor M (RimM) domain and a C2H2-type zinc finger domain. Both genes are widely conserved among the apicomplexans, and gene deletions led to the complete loss of virulence in mice, suggesting that TgGTPase and TgRimM are essential for the parasite adaptation to the *in vivo* environment. Overall, our current study provides a clue for the research community to estimate the *in vivo* essentiality of proteins with previously unknown localizations.

## RESULTS

### *In vivo* CRISPR screening targeting proteins of unassigned subcellular localization

To identify genes necessary for *Toxoplasma* survival in mice, we utilized a previously established *in vivo* CRISPR screen platform using type I strain and C57BL/6 mice (14). We newly generated two gRNA libraries (Unassigned_1 and Unassigned_2 libraries), each targeting 295 genes classified as unassigned localization in hyperLOPIT (7) and control genes for *in vitro* and *in vivo* selection. Among approximately 1200 hyperLOPIT-unassigned proteins, we excluded genes with previously published *in vitro* fitness scores in human foreskin fibroblasts (HFF) less than −1.5 to reduce the size of the gRNA library (10). Transfected RHΔhxgprt parasites were selected with pyrimethamine in Vero cells for four passages to generate a pooled mutant population (*in vitro* sample). The parasite mutant pools were used to infect C57BL/6 mice with 10^7^ parasites each by injection into the footpad. After seven days post infection, parasites were recovered from the spleen and expanded in Vero cells for one passage (*in vivo* sample). The gRNA sequences were amplified by PCR from the input library and the genomic DNA extracted from the parasites of *in vitro* and *in vivo* samples. The gRNAs were sequenced by next-generation sequencing to determine their relative abundance.

We calculated *in vitro* and *in vivo* fitness scores of each gene as the average log_2_ fold change of guide abundances between conditions. We analyzed the screen results from both sublibraries to assess the reproducibility **(Tables S1 and S2)**. The *in vitro* essential and dispensable controls were successfully separated, with lower scores for the essential genes and higher scores for the dispensable genes (**Figure S1A**). The correlations between our *in vitro* fitness scores and the genome-wide *in vitro* fitness scores in HFFs were high (r = 0.78 and 0.76, respectively) (**Figure S1B**) (10). High reproducibility of *in vivo* fitness scores was observed between independent infections in mice (r = 0.75±0.05 and 0.68±0.07, respectively) (**Figure S1C**). These data demonstrated that our *in vitro* and *in vivo* CRISPR screens using Unassigned_1 and Unassigned_2 libraries were highly reproducible.

To identify genes that confer parasite *in vivo* fitness during infection in mice, we compared *in vitro* and *in vivo* fitness scores (**Figure 1A, Tables S1 and S2**). To rank the candidate genes, we calculated the distance of each gene from the regression line (**Figure 1B**). As reported previously (14), the *in vivo* essential controls ROP18 and GRA23 showed negative *in vivo* fitness scores and ranked highly (**Figure 1B**). This analysis highlighted both previously identified and unidentified genes. For instance, the top hits from Unassigned_1 library contain hypothetical protein (TGGT1_211850), thioredoxin (Txn: TGGT1_224060), NUDIX hydrolase (Nudix: TGGT1_282190), CAAX metallo endopeptidase (Caax: TGGT1_221170), vacuolar iron transporter (VIT: TGGT1_266800), DNA damage inducible protein 1 (DDI1: TGGT1_304680), TgGTPase (TGGT1_277840), GLT2 (TGGT1_239752), apical annuli methyltransferase (AAMT: TGGT1_310070), and mRNA cleavage factor family protein (mRNACF: TGGT1_221190). It has been reported that VIT mainly localizes to the plant-like vacuolar compartment (PLVAC) and regulates parasite iron metabolism (16). DDI1 is one of the components of the parasite ubiquitin-proteasome system and is localized in the cytoplasm and nucleus (17). TgGTPase was identified as an interactor of the Nd complex, which facilitates rhoptry exocytosis (18). GLT2 is a glucosyltransferase and required for parasite disaccharide metabolism (19). AAMT was identified as an apical annuli component (20). The top hits from Unassigned_2 library also contain several hypothetical proteins (TGGT1_300220, TGGT1_312840, TGGT1_277870, TGGT1_250790, TGGT1_221640, and TGGT1_273905), ribosome maturation factor RimM domain-containing protein (TgRimM: TGGT1_321310), TBC6 (TGGT1_237280), DnaJ domain-containing protein (Dnaj: TGGT1_203380), CPSF A subunit region protein (CPSF: TGGT1_ 267710), and rRNA-processing protein EFG1(EFG1: TGGT1_240660). Although TGGT1_300220 was also shown to be one of the top hits in another *in vivo* CRISPR screen (15), no further investigation was performed. TBC6 is one of the TBC-domain containing proteins and is localized in the ER or cytoplasmic vesicles (21). TGGT1_273905 localizes to the cytoplasm (22). Although some candidates (VIT and DDI1) were previously reported to be required for parasite virulence (16, 17), none of the other top hits are assessed regarding virulence so far.

**Figure 1.**
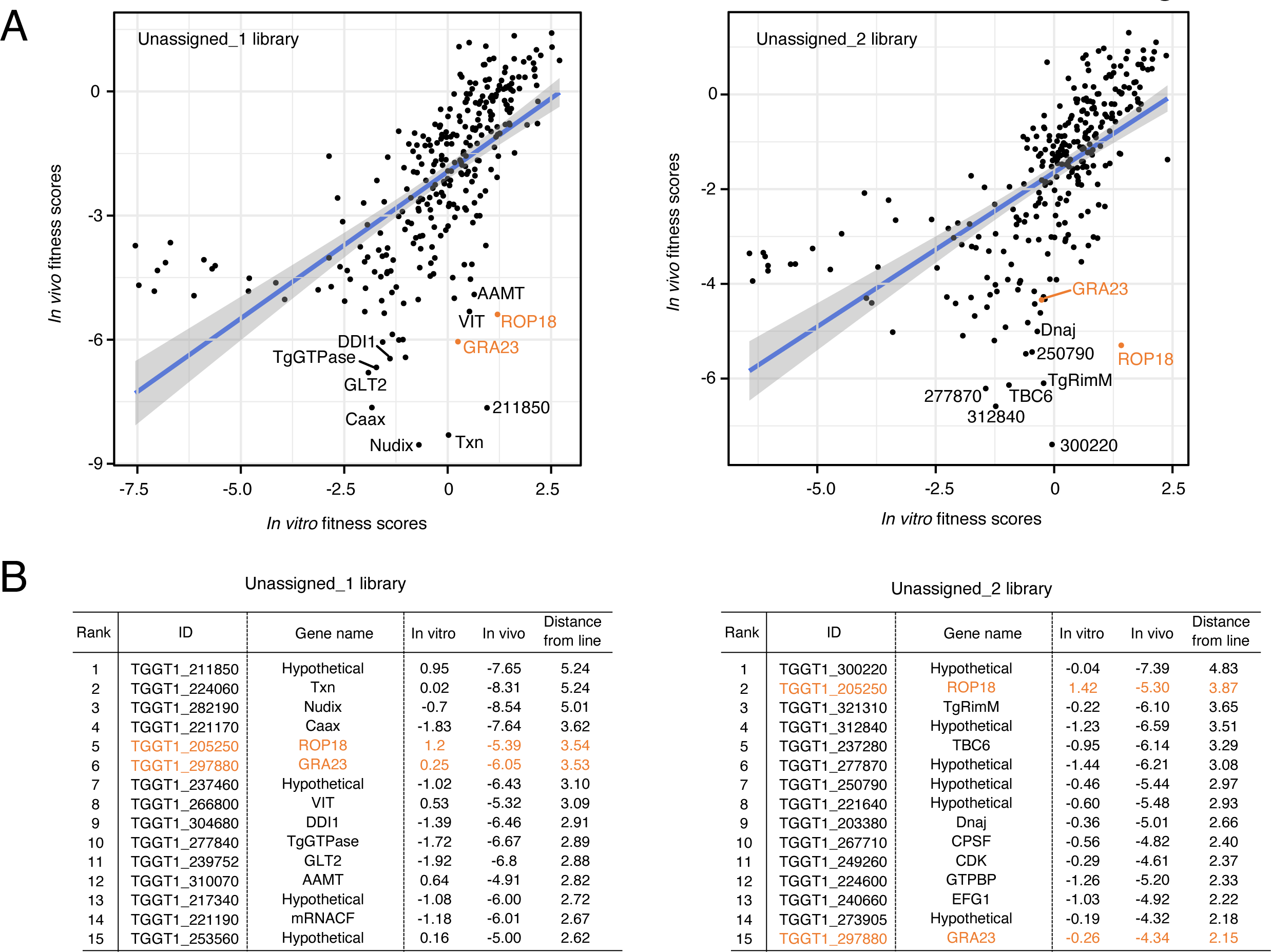
*In vivo* CRISPR screening targeting proteins with hyperLOPIT-unassigned localization. (A) Scatterplots showing in vitro and in vivo fitness scores from Unnassigned_1 (left) and Unassigned_2 (right) libraries, respectively. ROP18 and GRA23 are labeled in orange as control genes. (B) Ranking tables for top 15 *in vivo* fitness-conferring genes in each library ordered by the distance from the regression line.

### Identification of novel genes that contribute to parasite virulence in mice

We generated individual knockout strains to examine whether these top-hit genes affect parasite virulence (**Figures S2A and S2B**). WT mice were locally infected with these mutant strains into the footpad, and the survival rates were monitored. We chose footpad infection rather than intraperitoneal infection because our previous study revealed that the former is more suitable for assessing virulence in using RH strain (14) (**Figures 2A and 2B**). Among the candidates from Unassigned_1 library, all mice survived in the infection of ΔTGGT1_211850, ΔDDI1, ΔTgGTPase, or ΔAAMT parasites. Other tested mutants also showed reduced virulence (ΔGLT2, ΔCaax, ΔmRNACF, ΔTxn, ΔNudix, and ΔVIT). Among the candidates from Unassigned_2 library, all mice survived in the infection of ΔTGGT1_300220, ΔTgRimM, ΔEFG1, ΔCPSF, or ΔTGGT1_277870 parasites. ΔTGGT1_250790, ΔDnaj, ΔTGGT1_312840, or ΔTGGT1_273905 parasites showed reduced virulence. WT mice infected with ΔTBC6, and ΔTGGT1_221640 succumbed in a time course similar to those infected with WT parasites. Collectively, these data demonstrated that our *in vivo* CRISPR screens targeting hyperLOPIT-unassigned proteins highlighted known and novel *in vivo* fitness-conferring genes that affect virulence.

**Figure 2.**
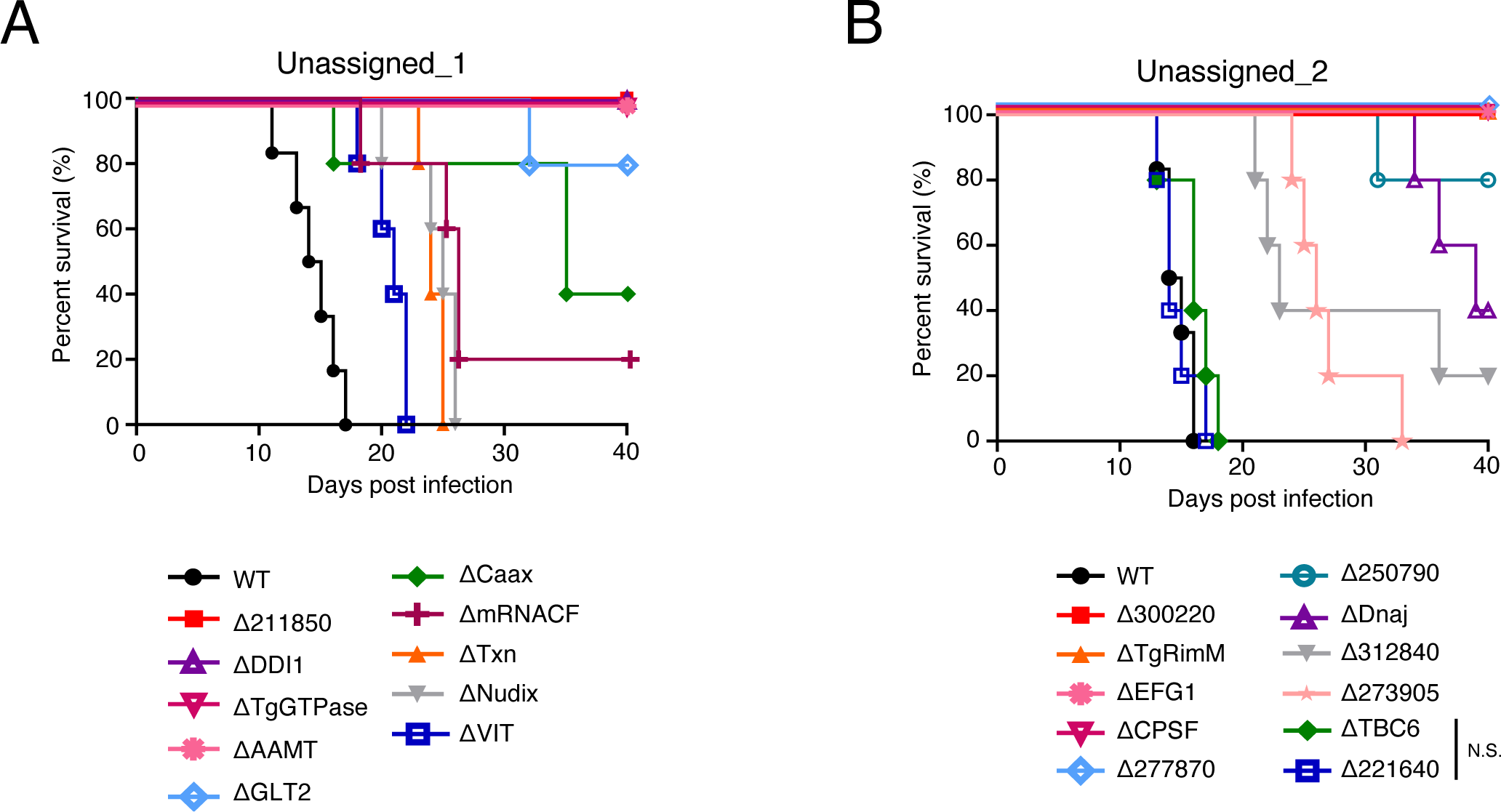
Validation of candidate genes that affect parasite virulence. (A) C57BL/6 mice were inoculated into the footpad with 10^3^ tachyzoites of the indicated knockout strains from the unassigned_1 library. WT (n = 6), ΔTGGT1_211850 (n = 5), ΔDDI1 (n = 5), ΔTgGTPase (n = 5), ΔAAMT (n = 5), ΔGLT2 (n = 5), ΔCaax (n = 5), ΔmRNACF (n = 5), ΔTxn (n = 5), ΔNudix (n = 5), ΔVIT (n = 5). (B) C57BL/6 mice were inoculated into the footpad with 10^3^ tachyzoites of the indicated knockout strains from the unassigned_2 library. WT (n = 6), ΔTGGT1_300220 (n = 5), ΔTgRimM (n = 5), ΔEFG1 (n = 5), ΔCPSF (n = 5), ΔTGGT1_277870 (n = 5), ΔTGGT1_250790 (n = 5), ΔDnaj (n = 5), ΔTGGT1_312840 (n = 5), ΔTGGT1_273905 (n = 5), ΔTBC6 (n = 5), ΔTGGT1_221640 (n = 5). N.S., not significant (log-rank test).

### TgGTPase is a widely conserved GTPase that contributes to virulence

Among the top-ranking candidates from Unassigned_1 library, we chose to characterize TgGTPase further (**Figure 3A and Tables S1**). TgGTPase is a GTPase that is broadly conserved across the phylum Apicomplexa (**Figures 3B and 3C**). To study the function of TgGTPase, we complemented the ΔTgGTPase strain with C-terminal HA-tagged TgGTPase expressed from the tubulin promoter that we will refer to as “ΔTgGTPase::TgGTPase-HA” (**Figure 3D**). Western blot showed the band of TgGTPase-HA at around 40 kDa (**Figure 3E**). The HA-tagged TgGTPase showed a cytoplasmic localization by IFA (**Figure 3F**). To validate the essentiality of TgGTPase on pathogenesis, we infected C57BL/6 mice intraperitoneally with 10^3^ tachyzoites of the WT, ΔTgGTPase, and ΔTgGTPase::TgGTPase-HA parasites and assessed mouse survival (**Figure 3G**). All the mice intraperitoneally infected with ΔTgGTPase parasites survived. By contrast, ΔTgGTPase::TgGTPase-HA parasites fully restored virulence as well as WT parasites. Taken together, these results suggest that TgGTPase is a cytoplasmic protein essential for virulence.

**Figure 3.**
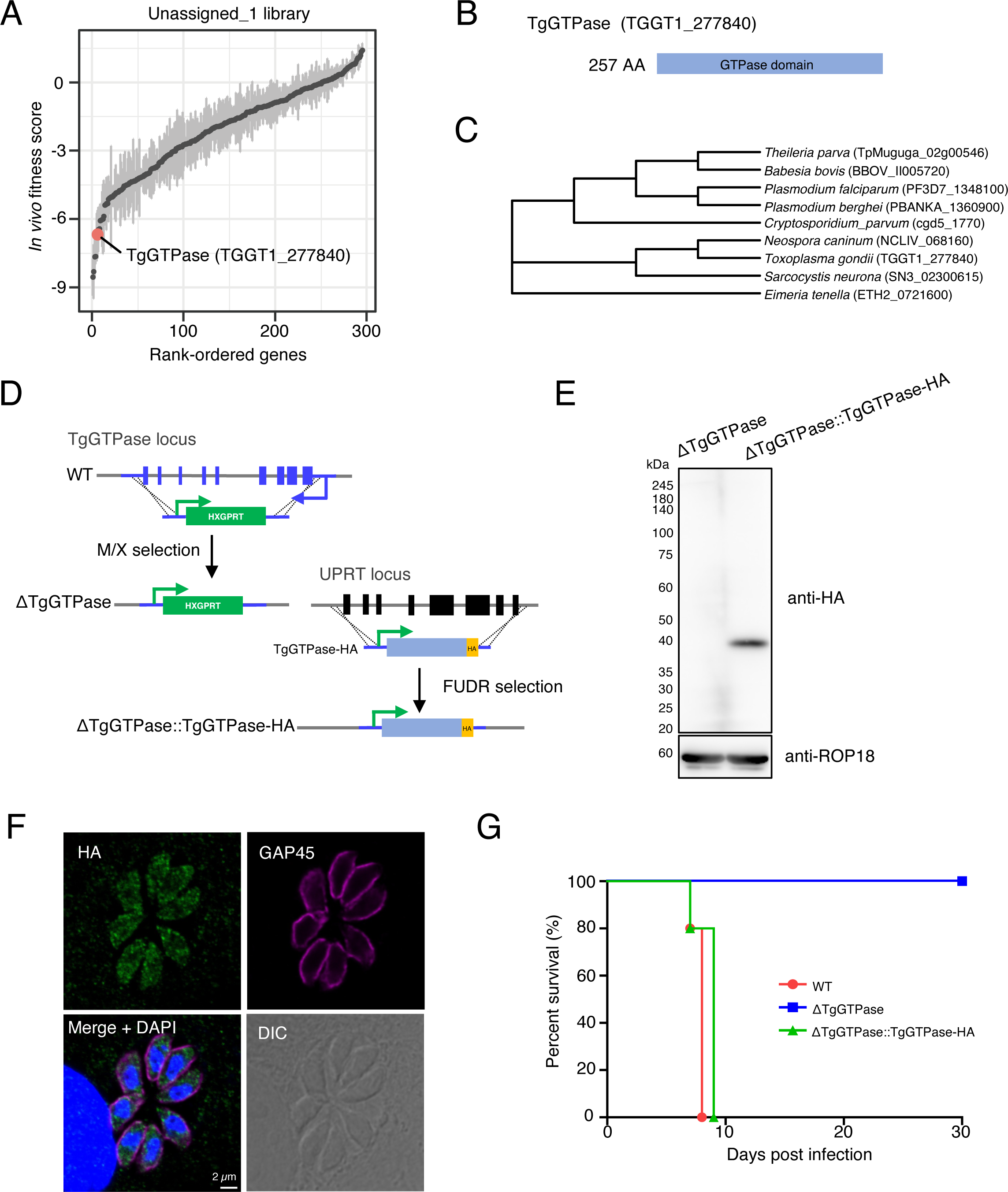
TgGTPase is a cytosolic GTPase that contributes to parasite virulence. (A) Rank-ordered plot of genes from Unassigned_1 library. (B) Schematic of putative domains in TgGTPase. (C) Phylogenetic tree of selected TgGTPase homologs across the apicomplexan phylum. (D) Schematic of TgGTPase knockout and complementation of TgGTPase-HA. (E) Western blot analysis of TgGTPase-HA. (F) Immunofluorescence images of ΔTgGTPase::TgGTPase-HA showing cytosolic localization of TgGTPase-HA (green). GAP45 (magenta) stains the inner membrane complex of the parasites. (G) Survival curves of C57BL/6 mice intraperitoneally infected with 10^3^ tachyzoites of WT (n = 5), ΔTgGTPase (n = 5), and ΔTgGTPase::TgGTPase-HA (n = 5). Data are representative of three independent experiments (E and F).

### TgRimM is a novel apicoplast-resident protein possessing C2H2 zinc finger that contributes to virulence in mice

Among the top-ranking candidates from Unassigned_2 library, we chose to characterize TgRimM further (**Figure 4A and Tables S2**). TgRimM possesses a signal peptide, a ribosome maturation factor RimM domain, and a single C2H2-type zinc finger domain (**Figure 4B**) (23, 24). We searched for TgRimM homologs and found that they are broadly conserved across the phylum Apicomplexa except for *Cryptosporidium* spp. (**Figure 4C**). To characterize TgRimM, we complemented the ΔTgRimM strain with C-terminal HA-tagged TgRimM expressed from the tubulin promoter that we will refer to as “ΔTgRimM::TgRimM-HA” (**Figure 4D**). The HA-tagged TgRimM showed an apicoplast localization co-stained with streptavidin (an apicoplast marker) by IFA (**Figure 4E**). Western blot against TgRimM-HA showed the main band at around 75 kDa, probably a processed form of TgRimM (**Figure 4F**). To check the effect of TgRimM on virulence, we infected C57BL/6 mice intraperitoneally with 10^3^ tachyzoites of the WT, ΔTgRimM, and ΔTgRimM::TgRimM-HA parasites and assessed the mouse survival. All the mice intraperitoneally infected with ΔTgRimM parasites survived. In contrast, ΔTgRimM::TgRimM-HA parasites restored virulence (**Figure 4G**). Next, we assessed the importance of the C2H2-type zinc finger domain in TgRimM. We generated a complemented strain lacking the C2H2 zinc finger (ΔTgRimM::TgRimM^ΔZF^-HA) (**Figure 4C**). Deleting the zinc finger domain did not alter the apicoplast localization of TgRimM^ΔZF^-HA (**Figure S3**). However, ΔTgRimM::TgRimM^ΔZF^-HA could not restore virulence (**Figure 4G**). Thus, these results suggest that TgRimM is essential for virulence in mice and that the C2H2 zinc finger is indispensable for TgRimM function in the apicoplast.

**Figure 4.**
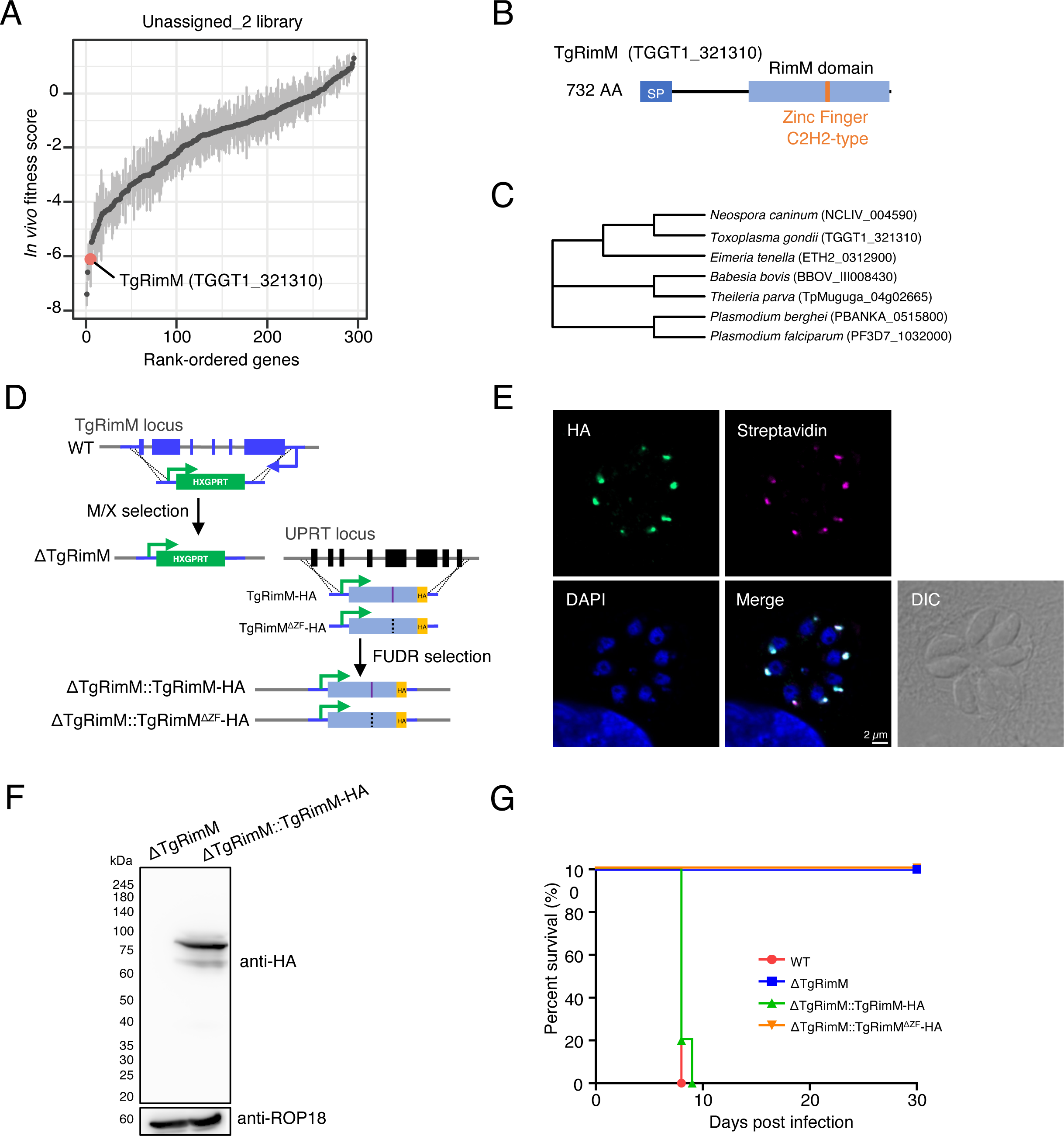
TgRimM is an apicoplast-resident protein that is essential for parasite virulence. (A) Rank-ordered plot of genes from Unassigned_2 library. (B) Schematic of putative domains in TgRimM. SP is predicted by SignalP 6.0. (C) Phylogenetic tree of selected TgRimM homologs across the apicomplexan phylum. (D) Schematic of TgRimM knockout and complementation of wild-type and mutant TgRimM. (E) Immunofluorescence images of ΔTgRimM::TgRimM-HA showing TgRimM-HA (green) is co-stained with streptavidin (magenta), an apicoplast marker. (F) Western blot analysis of TgRimM-HA. (G) Survival curves of C57BL/6 mice intraperitoneally infected with 10^3^ tachyzoites of WT (n = 5), ΔTgRimM (n = 5), ΔTgRimM::TgRimM-HA (n = 5), or ΔTgRimM::TgRimM^ΔZF^-HA (n = 5). Data are representative of three independent experiments (E and F).

## DISCUSSION

Despite much effort from the research community, many apicomplexan proteins still need to be annotated owing to their uniqueness and absence in other model organisms. The CRISPR screens enabled the researchers to investigate hundreds to thousands of genes in *Toxoplasma* at once *in vitro* culture or *in vivo* infection to identify essential genes for parasite growth, metabolism, drug resistance, differentiation, host-pathogen interaction, and virulence (10–15, 25–33). Most *in vivo* CRISPR screens in *Toxoplasma* have been focused on identifying secreted virulence effectors such as rhoptry bulb proteins (ROPs) and dense granule proteins (GRAs) (11–13). However, recent studies are in progress to search for non-secreted virulence factors by *in vivo* CRISPR screens (14, 15). In this study, we focused on the hyperLOPIT-unassigned proteins and performed *in vivo* CRISPR screens to identify novel virulence factors in *Toxoplasma*. As a result, we identified many candidate genes that were required for the parasite *in vivo* fitness in mice and further characterized TgGTPase and TgRimM among them.

TgGTPase was originally reported as an interactor of the Nd complex, which plays an important role in rhoptry secretion (18). However, considering that our study revealed its distribution in the cytosol, TgGTPase might have broader functions than rhoptry secretion. TgGTPase homologs are widely conserved across the phylum Apicomplexa, suggesting that this GTPase is functionally important across the species in the phylum. In the transposon mutagenesis screen in *Plasmodium falciparum* (34), PfGTPase (PF3D7_1348100) possesses lower mutagenesis scores (Mutagenesis Index Score = 0.316 and Mutagenesis Fitness Score = −2.437), suggesting that PfGTPase is essential for optimal growth of asexual blood-stages *in vitro*. Future studies on TgGTPase homologs in *Plasmodium* or *Cryptosporidium* will reveal their conserved function in the phylum.

We identified TgRimM as an apicoplast-resident protein. A homolog in *P. falciparum* (PfRimM: PF3D7_1032000) is also predicted to be localized in the apicoplast by signal peptide prediction (35). In the transposon mutagenesis screen in *P. falciparum* (34), PfRimM also possesses very low mutagenesis scores (Mutagenesis Index Score = 0.158 and Mutagenesis Fitness Score = −2.8), indicating that PfRimM is essential for growth of asexual blood-stages *in vitro*. Assuming that RimM homologs in the phylum Apicomplexa (ApiRimM) are located in the apicoplast, the lack of RimM homologs in *Cryptosporidium* is reasonable because *Cryptosporidium* spp. are known to have lost the apicoplast (36–38). RimM domain-containing protein was reported to regulate 16S rRNA processing in bacteria (39). It is reported that a RimM domain-containing protein in *Arabidopsis thaliana* (AtRimM) is located in the chloroplasts and contributes to rRNA maturation and proteostasis (40). Considering that the apicoplast evolved from secondary endosymbiosis (41) and that the zinc finger domain in TgRimM is essential for its function in our study, ApiRimM might facilitate rRNA maturation and proteostasis in the apicoplast. Previous studies on *Toxoplasma* apicoplast have shown that apicoplast function is essential for the parasite *in vitro* survival and growth (42–44). Moreover, apicoplast-assigned proteins in hyperLOPIT showed a small number of *in vitro* dispensable proteins (7), also supporting its bare essentiality. In sharp contrast, other than TgRimM, we previously identified two additional apicoplast-resident proteins (EF-P: TGGT1_258380 and hypothetical protein: TGGT1_204350) that are dispensable for *in vitro* fitness but essential for *in vivo* fitness in CRISPR screens (14). The presence of this type of apicoplast proteins that are dispensable *in vitro* but indispensable *in vivo* might imply unknown characteristics of the apicoplast. Considering that the apicoplast is a potent drug target against toxoplasmosis and malaria (45), the presence of such apicoplast proteins that are dispensable *in vitro* but indispensable *in vivo* might expand the range of anti-toxoplasmosis and -malarial drug targets in the future. In combination with the database of hyperLOPIT, *in vivo* CRISPR screens succeeded in identifying such “new class” apicoplast proteins essential for virulence.

Limitations of our current study are that, although approximately 1,200 proteins are classified as unassigned subcellular localizations in hyperLOPIT, our sublibraries cover only half of them. Therefore, the remaining proteins still need to be addressed in the future. Also, although we identified previously unreported virulence genes in this study, their precise functions and localizations, other than TgGTPase and TgRimM, mostly remained unaddressed. Future works will aim to characterize these remaining proteins further.

In conclusion, our *in vivo* CRISPR screen platform provides a resource for the community to identify virulence genes encoding proteins with previously unknown localizations and functions.

## MATERIALS AND METHODS

### Toxoplasma strains

RHΔhxgprt (14), RHΔhxgprtΔku80 (46) and its derivatives of *Toxoplasma* were maintained in Vero cells and passaged every 3 days using RPMI (Nacalai Tesque) supplemented with 2 % heat-inactivated fetal bovine serum (FBS; JRH Bioscience), 100 U/ml penicillin and 0.1 mg/ml streptomycin (Nacalai Tesque) in incubators at 37°C and 5% CO_2_.

### Host cell culture

Vero cells were maintained in RPMI (Nacalai Tesque) supplemented with 10 % heat-inactivated FBS, 100 U/ml penicillin and 0.1 mg/ml streptomycin (Nacalai Tesque) in incubators at 37°C and 5% CO_2_. Human foreskin fibroblasts (HFFs) were maintained in DMEM (Nacalai Tesque) supplemented with 10% heat-inactivated FBS, 100 U/ml penicillin and 0.1 mg/ml streptomycin (Nacalai Tesque) in incubators at 37°C and 5% CO_2_.

### Mice

C57BL**/**6NCrSlc (C57BL/6N) mice were purchased from SLC. All experiments were conducted in 8-10-week-old female mice. All animal experiments were conducted with the approval of the Animal Research Committee of Research Institute for Microbial Diseases in Osaka University.

### *In vitro* and *in vivo* pooled CRISPR screens

The gRNA sequences of Unassigned_1 and Unassigned_2 sublibrary were selected from the genome-wide gRNA library (10). The selected gRNA sequences were cloned into the modified pU6-Universal vector by cloning a T2A, DHFR, T2A, and RFP in frame with Cas9, where the expression of gRNA and Cas9-T2A-DHFR-T2A-RFP cassettes was independently transcribed. The insertion of the selected gRNA sequences into the vector was performed by VectorBuilder. The gRNA library (200 μg) was linearized with NotI and transfected into approximately 1-2×10^8^ RHΔhxgprt parasites divided between four separate cuvettes. Then, transfected parasites were grown in 4×150-mm dishes with confluent Vero cell monolayers. Pyrimethamine (Sigma) was added 24 h post transfection. All the parasites were passaged every 3 days until Passage 3 without filtration. After 2 days (Passage 4), the parasites were syringe lysed, filtered, and counted for genomic DNA preparation or for mouse infection. For genomic DNA preparation, at least 1×10^8^ parasites were pelleted and stored at −80 °C. For mouse infection, the parasites were resuspended in PBS at a concentration of 2.5×10^8^ parasites/ml. Then, 1×10^7^ parasites in 40 μl PBS were injected into the footpad of each anesthetized mouse. Parasite viability was determined by plaque assay. At 7 days post infection, the spleens were collected and crushed by a plunger and passed through a cell strainer to make single cell suspensions. Then, the suspensions were pelleted and added to 2×150-mm dishes per spleen with confluent Vero cell monolayers. After 2-4 days, when the parasites completely lysed out, they were filtered and counted. At least 1×10^8^ parasites were pelleted and stored at −80 °C. Parasite genomic DNA was extracted using the DNeasy Blood and Tissue kit (Qiagen) according to the manufacturer’s instructions. Integrated gRNA sequences were PCR-amplified and barcoded with KOD FX Neo (TOYOBO) using Primer 1 and Primer 2 (**Table S3**). Genomic DNA (1μg) was used for the template. The resulting libraries were sequenced on a DNBSEQ-G400RS (MGI) using Primer 3 and Primer 4 (**Table S3)**.

### Bioinformatic analysis of the CRISPR screen

Following demultiplexing, gRNA sequencing reads were aligned to the gRNA library. The abundance of each gRNA was calculated and normalized to the total number of aligned reads (47). For *in vitro* analysis, the log_2_ fold-change between the P4 sample and the library was calculated for each gRNA. The fitness score for each gene was calculated as the mean log_2_ fold change for the top five scoring guides. For *in vivo* analysis, the log_2_ fold-change between each *in vivo* sample and the P4 sample was calculated for each gRNA as described above. The median fitness score across mouse replicates was used as the *in vivo* fitness score (14). For a given gene, gRNAs were compared using Wilcoxon rank sum test between P4 vs WT mice. The p values for each test were adjusted using the Benjamini-Hochberg method. The distance of each gene from the regression line was calculated as below.

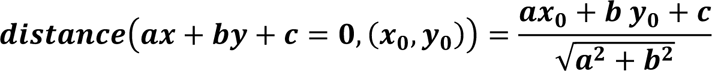

All analyses were conducted by R (v4.1.1) with package stats (v3.6.2) and visualized by ggplot2 (v3.4.0).

### Plasmid construction for knockout *Toxoplasma*

For construction of the CRISPR/Cas9 plasmids for targeting a gene of interest (GOI), two oligonucleotide primers (GOI_gRNA1_F and GOI_gRNA1_R, GOI_gRNA2_F and GOI_gRNA2_R) containing gRNA sequence were annealed and cloned into BsaI site of the pU6-Universal vector (Addgene #52694). To generate a construct for deleting the entire coding region of GOI, flanking regions of 60 bp of 5’ and 3’ outside the gRNAs were used to surround the genes. Forward and reverse primers with homology to the floxed HXGPRT cassette and to the 5’ and 3’ coding sequence of the GOI was used (GOI_homology_F and GOI_ homology _R). The primer sequences used for the genetic disruption are shown in **Table S3**.

### Generation of gene knockout *Toxoplasma*

RHΔhxgprtΔku80 were filtered and resuspended in Cytomix (10mM KPO_4_, 120mM KCl, 0.15 mM CaCl_2_, 5mM MgCl_2_, 25 mM HEPES, 2 mM EDTA). Parasites were mixed with 50 µg of each gRNA1 and gRNA2 CRISPR plasmids with the PCR-amplified targeting fragment for each GOI, and supplemented with 2 mM ATP, 5mM GSH. Parasites were electroporated by GENE PULSER II (Bio-Rad Laboratories). Transfected parasites were selected by 25 µg/ml mycophenolic acid (MPA) (Sigma) and 50 µg/ml Xanthine (Wako) to obtain stably resistant clones. Then, parasites were subjected to limiting dilution in 96-well plates to isolate single clones. To confirm the disruption of the gene, we analyzed the mRNA expression by quantitative RT-PCR.

### Complementation of TgGTPase and TgRimM

To complement the ΔTgGTPase and ΔTgRimM parasites, TgGTPase, TgRimM, and TgRimM^ΔZF^ cDNA was generated by PCR amplification from cDNA of the RH strain. The PCR products were inserted into the pUPRT plasmid vector, which possesses cDNA of interest with SPOT and HA epitope tag driven by the tubulin promoter. Indicated parasites were transfected with the pUPRT complementation vector and gRNA targeting the UPRT locus, then selected by FUDR (Wako) (48). Isolated clones were examined for protein expression by western blotting and IFA.

### TgGTPase and TgRimM phylogenetic analysis

ToxoDB (23), VEuPathDB (49) and BLAST were searched for TgGTPase or TgRimM homologs across the phylum Apicomplexa. Clustal Omega was used to align obtained homologs (50). Phylogenetic trees were visualized by ggtree (v3.2.1) (51).

### Immunofluorescence assay

HFFs were grown on coverslips, infected with parasites for 24-30 h, and fixed in PBS containing 3.7% paraformaldehyde (PFA) for 10 min at room temperature. Cells were permeabilized with PBS containing 0.1% Triton-X for 10 min and then blocked with in PBS containing 3% BSA for 1 h at room temperature. Then, the coverslips were incubated with the primary antibodies for 1 h, followed by incubation with appropriate secondary antibodies, streptavidin and DAPI for 30 min. The coverslips were mounted using PermaFluor (Thermo Scientific). Images were acquired by confocal laser microscopy (Olympus FV3000 IX83). Primary antibodies used were mouse anti-HA.11 (BioLegend, 901514), rabbit anti-GAP45 (52), and Alexa Fluor 594 streptavidin (Invitrogen, S11227).

### Assessment of *in vivo* virulence in mice

Mice were infected with 10^3^ tachyzoites in 200 µl (intra-peritoneum) or 40 µl (intra-footpad) of PBS. Parasite viability was determined by plaque assay. The mouse health condition and survival were monitored daily until 30 days (intra-peritoneum) or 40 days (intra-footpad) post infection, respectively.

### Quantitative RT-PCR

Total RNA was extracted by RNeasy kit (QIAGEN), and cDNA was synthesized by Verso reverse transcription (Thermo Fisher Scientic) according to the manufacturer’s instructions. Quantitative RT-PCR was performed with a CFX Connect real-time PCR system (Bio-Rad Laboratories) and a Go-Taq real-time PCR system (Promega). The data were analyzed by the ΔΔCT method and normalized to ACT1 in each sample. The primer sequences are listed in **Table S3**.

### Western blotting

Cells were lysed in lysis buffer (1% NP-40, 150 mM NaCl, 20 mM Tris-HCl, pH 7.5) containing a protease inhibitor cocktail (Nacalai Tesque). The cell lysates were separated by SDS-PAGE and transferred to polyvinylidene difluoride membranes (Immobilon-P; Millipore). Membranes were blocked in 5% skim milk in 0.05% Tween-20 in PBS for 1 h at room temperature, followed by incubation with primary antibodies for 1 h at room temperature. Blots were then incubated with appropriate secondary antibodies for 1 h at room temperature. Primary antibodies used were mouse anti-HA (MBL, M180-7) and rabbit anti-ROP18 (this study).

### Generation of custom anti-ROP18 antibody

Custom anti-ROP18 antibody (rabbit polyclonal) was generated against a synthetic C-terminal peptide of ROP18 (AQNFEQQEHLHTE). The epitope identification, peptide synthesis, rabbit immunization and serum collection were conducted by Cosmo Bio. The specificity of anti-ROP18 antibody was validated for western blotting.

### Quantification and statistical analysis

Information about the number of biological replicates and the type of statistical tests used can be found in the figure legends. All statistical analyses except for the survival rates were performed using R (4.1.1, https://www.r-project.org/). For correlation analysis, Pearson’s correlation was used. Data with p values < 0.05 were considered statistically significant. The statistical analysis of survival rates was performed by the log-rank test using the GraphPad Prism9 software.

## Data availability

The CRISPR screen data of two sublibraries have been deposited to the NCBI GEO. The GEO accession numbers are **GSE253884** and **GSE253885**. Any additional information required to reanalyze the data reported in this paper is available from the corresponding author upon request.

## ACKNOWLEGDEMENT

We thank M. Enomoto and N. Yamagishi (Osaka University) for secretarial and technical assistance. We acknowledge the NGS core facility at the Research Institute for Microbial Diseases of Osaka University for the sequencing. We thank Dr. D. Soldati-Favre for anti-GAP45 antibody. This study was supported by Fusion Oriented Research for Disruptive Science and Technology (JPMJFR206D), and Moonshot research & development (JPMJMS2025) from Japan Science and Technology Agency, the Research Program on Emerging and Re-emerging Infectious Diseases (JP23fk0108682) from the Agency for Medical Research and Development (AMED), a Grant-in-Aid for Transformative Research Area (B) (Establishment of PLAMP as a new concept to determine self and nonself for obligatory intracellular pathogens; 20B304), for Scientific Research (A) (19H00970), for JSPS Research Fellow (23KJ1469) from the Ministry of Education, Culture, Sports, Science and Technology, program from Joint Usage and Joint Research Programs of the Institute of Advanced Medical Sciences, Tokushima University, Takeda Science Foundation, Mochida Memorial Foundation, Astellas Foundation for Research on Metabolic Disorders, Naito Foundation, the Chemo-Sero-Therapeutic Research Institute, Research Foundation for Microbial Diseases of Osaka University, BIKEN Taniguchi Scholarship, and Joint Research Program of Research Center for Global and Local Infectious Diseases of Oita University (2021B06).

## AUTHOR CONTRIBUTIONS

Conceptualization, Y.T. and M.Y.; Methodology, Y.T. and M.Y.; Software and Formal Analysis, Y.T.; Investigation, Y.T. and M.Y.; Writing – Original Draft, Y.T. and M.Y.; Writing – Review & Editing, Y.T., M.S. and M.Y.; Funding Acquisition, M.Y.; Supervision, M.Y.

## SUPPLEMENTAL MATERIAL

**Supplementary Figure 1. Assessing the reproducibility of CRISPR screens.**

**Related to Figure 1.**

(A) Rank-ordered plots for *in vitro* fitness scores of 4^th^ passage.

(B) Correlation between our *in vitro* fitness scores and the *in vitro* fitness scores in HFF.

(C) Overlay of *in vivo* fitness scores for each mouse. Pearson’s correlation coefficients are shown as mean ± SD.

**Supplementary Figure 2. Generating gene knockout parasites.**

**Related to Figure 2.**

(A) Schematic of gene knockout strategy.

(B) Quantitative RT-PCR validations for indicated parasites.

**Supplementary Figure 3. TgRimMΔZF-HA localized in the apicoplast.**

**Related to Figure 4.**

Immunofluorescence images of ΔTgRimM::TgRimM^ΔZF^-HA showing TgRimM^ΔZF^-HA (green) is co-stained with streptavidin (magenta), an apicoplast marker. Data are representative of three independent experiments.

**Table S1.**

Summary of *in vivo* CRISPR screen using Unassigned_1 library, gRNA sequences, raw count data, and *in vivo* fitness scores for each mouse.

**Table S2.**

Summary of *in vivo* CRISPR screen using Unassigned_2 library, gRNA sequences, raw count data, and *in vivo* fitness scores for each mouse.

**Table S3.**

Lists of the primers used in this study.

## REFERENCES

1. Montoya JG, Liesenfeld O. 2004. Toxoplasmosis. Lancet 363:1965–76.

2. Kim K, Weiss LM. 2004. Toxoplasma gondii: the model apicomplexan. Int J Parasitol 34:423–32.

3. Lebrun M, Carruthers VB, Cesbron-Delauw M-F. 2020. Toxoplasma secretory proteins and their roles in parasite cell cycle and infection, p 607–704, Toxoplasma gondii. Elsevier.

4. Kloehn J, Lacour CE, Soldati-Favre D. 2021. The metabolic pathways and transporters of the plastid organelle in Apicomplexa. Curr Opin Microbiol 63:250–258.

5. Gambarotto D, Zwettler FU, Le Guennec M, Schmidt-Cernohorska M, Fortun D, Borgers S, Heine J, Schloetel JG, Reuss M, Unser M, Boyden ES, Sauer M, Hamel V, Guichard P. 2019. Imaging cellular ultrastructures using expansion microscopy (U-ExM). Nat Methods 16:71–74.

6. Tosetti N, Dos Santos Pacheco N, Bertiaux E, Maco B, Bournonville L, Hamel V, Guichard P, Soldati-Favre D. 2020. Essential function of the alveolin network in the subpellicular microtubules and conoid assembly in Toxoplasma gondii. Elife 9.

7. Barylyuk K, Koreny L, Ke H, Butterworth S, Crook OM, Lassadi I, Gupta V, Tromer E, Mourier T, Stevens TJ, Breckels LM, Pain A, Lilley KS, Waller RF. 2020. A Comprehensive Subcellular Atlas of the Toxoplasma Proteome via hyperLOPIT Provides Spatial Context for Protein Functions. Cell Host Microbe 28:752–766 e9.

8. Guerin A, Strelau KM, Barylyuk K, Wallbank BA, Berry L, Crook OM, Lilley KS, Waller RF, Striepen B. 2023. Cryptosporidium uses multiple distinct secretory organelles to interact with and modify its host cell. Cell Host Microbe 31:650–664 e6.

9. Christoforou A, Mulvey CM, Breckels LM, Geladaki A, Hurrell T, Hayward PC, Naake T, Gatto L, Viner R, Martinez Arias A, Lilley KS. 2016. A draft map of the mouse pluripotent stem cell spatial proteome. Nat Commun 7:8992.

10. Sidik SM, Huet D, Ganesan SM, Huynh MH, Wang T, Nasamu AS, Thiru P, Saeij JPJ, Carruthers VB, Niles JC, Lourido S. 2016. A Genome-wide CRISPR Screen in Toxoplasma Identifies Essential Apicomplexan Genes. Cell 166:1423–1435 e12.

11. Young J, Dominicus C, Wagener J, Butterworth S, Ye X, Kelly G, Ordan M, Saunders B, Instrell R, Howell M, Stewart A, Treeck M. 2019. A CRISPR platform for targeted in vivo screens identifies Toxoplasma gondii virulence factors in mice. Nat Commun 10:3963.

12. Sangare LO, Olafsson EB, Wang Y, Yang N, Julien L, Camejo A, Pesavento P, Sidik SM, Lourido S, Barragan A, Saeij JPJ. 2019. In Vivo CRISPR Screen Identifies TgWIP as a Toxoplasma Modulator of Dendritic Cell Migration. Cell Host Microbe 26:478–492 e8.

13. Butterworth S, Torelli F, Lockyer EJ, Wagener J, Song OR, Broncel M, Russell MRG, Moreira-Souza ACA, Young JC, Treeck M. 2022. Toxoplasma gondii virulence factor ROP1 reduces parasite susceptibility to murine and human innate immune restriction. PLoS Pathog 18:e1011021.

14. Tachibana Y, Hashizaki E, Sasai M, Yamamoto M. 2023. Host genetics highlights IFN-gamma-dependent Toxoplasma genes encoding secreted and non-secreted virulence factors in in vivo CRISPR screens. Cell Rep 42:112592.

15. Giuliano CJ, Wei KJ, Harling FM, Waldman BS, Farringer MA, Boydston EA, Lan TCT, Thomas RW, Herneisen AL, Sanderlin AG, Coppens I, Dvorin JD, Lourido S. 2023. Functional profiling of the Toxoplasma genome during acute mouse infection. bioRxiv doi:10.1101/2023.03.05.531216.

16. Aghabi D, Sloan M, Gill G, Hartmann E, Antipova O, Dou Z, Guerra AJ, Carruthers VB, Harding CR. 2023. The vacuolar iron transporter mediates iron detoxification in Toxoplasma gondii. Nat Commun 14:3659.

17. Zhang H, Liu J, Ying Z, Li S, Wu Y, Liu Q. 2020. Toxoplasma gondii UBL-UBA shuttle proteins contribute to the degradation of ubiquitinylated proteins and are important for synchronous cell division and virulence. FASEB J 34:13711–13725.

18. Aquilini E, Cova MM, Mageswaran SK, Dos Santos Pacheco N, Sparvoli D, Penarete-Vargas DM, Najm R, Graindorge A, Suarez C, Maynadier M, Berry-Sterkers L, Urbach S, Fahy PR, Guerin AN, Striepen B, Dubremetz JF, Chang YW, Turkewitz AP, Lebrun M. 2021. An Alveolata secretory machinery adapted to parasite host cell invasion. Nat Microbiol 6:425-434.

19. Gas-Pascual E, Ichikawa HT, Sheikh MO, Serji MI, Deng B, Mandalasi M, Bandini G, Samuelson J, Wells L, West CM. 2019. CRISPR/Cas9 and glycomics tools for Toxoplasma glycobiology. J Biol Chem 294:1104–1125.

20. Engelberg K, Chen CT, Bechtel T, Sanchez Guzman V, Drozda AA, Chavan S, Weerapana E, Gubbels MJ. 2020. The apical annuli of Toxoplasma gondii are composed of coiled-coil and signalling proteins embedded in the inner membrane complex sutures. Cell Microbiol 22:e13112.

21. Quan JJ, Nikolov LA, Sha J, Wohlschlegel JA, Bradley PJ. 2023. Toxoplasma gondii encodes an array of TBC-domain containing proteins including an essential regulator that targets Rab2 in the secretory pathway. bioRxiv doi:10.1101/2023.05.28.542599.

22. Roumegous C, Abou Hammoud A, Fuster D, Dupuy JW, Blancard C, Salin B, Robinson DR, Renesto P, Tardieux I, Frenal K. 2022. Identification of new components of the basal pole of Toxoplasma gondii provides novel insights into its molecular organization and functions. Front Cell Infect Microbiol 12:1010038.

23. Harb OS, Roos DS. 2020. ToxoDB: Functional Genomics Resource for Toxoplasma and Related Organisms. Methods Mol Biol 2071:27–47.

24. Teufel F, Almagro Armenteros JJ, Johansen AR, Gislason MH, Pihl SI, Tsirigos KD, Winther O, Brunak S, von Heijne G, Nielsen H. 2022. SignalP 6.0 predicts all five types of signal peptides using protein language models. Nat Biotechnol 40:1023–1025.

25. Butterworth S, Kordova K, Chandrasekaran S, Thomas KK, Torelli F, Lockyer EJ, Edwards A, Goldstone R, Koshy AA, Treeck M. 2023. High-throughput identification of Toxoplasma gondii effector proteins that target host cell transcription. Cell Host Microbe 31:1748–1762 e8.

26. Chen Y, Liu Q, Xue JX, Zhang MY, Geng XL, Wang Q, Jiang W. 2021. Genome-Wide CRISPR/Cas9 Screen Identifies New Genes Critical for Defense Against Oxidant Stress in Toxoplasma gondii. Front Microbiol 12:670705.

27. Harding CR, Sidik SM, Petrova B, Gnadig NF, Okombo J, Herneisen AL, Ward KE, Markus BM, Boydston EA, Fidock DA, Lourido S. 2020. Genetic screens reveal a central role for heme metabolism in artemisinin susceptibility. Nat Commun 11:4813.

28. Krishnamurthy S, Maru P, Wang Y, Bitew MA, Mukhopadhyay D, Yamaryo-Botte Y, Paredes-Santos TC, Sangare LO, Swale C, Botte CY, Saeij JPJ. 2023. CRISPR Screens Identify Toxoplasma Genes That Determine Parasite Fitness in Interferon Gamma-Stimulated Human Cells. mBio 14:e0006023.

29. Krishnan A, Kloehn J, Lunghi M, Chiappino-Pepe A, Waldman BS, Nicolas D, Varesio E, Hehl A, Lourido S, Hatzimanikatis V, Soldati-Favre D. 2020. Functional and Computational Genomics Reveal Unprecedented Flexibility in Stage-Specific Toxoplasma Metabolism. Cell Host Microbe 27:290–306 e11.

30. Lockyer EJ, Torelli F, Butterworth S, Song OR, Howell S, Weston A, East P, Treeck M. 2023. A heterotrimeric complex of Toxoplasma proteins promotes parasite survival in interferon gamma-stimulated human cells. PLoS Biol 21:e3002202.

31. Paredes-Santos TC, Bitew MA, Swale C, Rodriguez F, Krishnamurthy S, Wang Y, Maru P, Sangare LO, Saeij JPJ. 2023. Genome-wide CRISPR screen identifies genes synthetically lethal with GRA17, a nutrient channel encoding gene in Toxoplasma. PLoS Pathog 19:e1011543.

32. Waldman BS, Schwarz D, Wadsworth MH, 2nd, Saeij JP, Shalek AK, Lourido S. 2020. Identification of a Master Regulator of Differentiation in Toxoplasma. Cell 180:359-372 e16.

33. Wang Y, Sangare LO, Paredes-Santos TC, Hassan MA, Krishnamurthy S, Furuta AM, Markus BM, Lourido S, Saeij JPJ. 2020. Genome-wide screens identify Toxoplasma gondii determinants of parasite fitness in IFNgamma-activated murine macrophages. Nat Commun 11:5258.

34. Zhang M, Wang C, Otto TD, Oberstaller J, Liao X, Adapa SR, Udenze K, Bronner IF, Casandra D, Mayho M, Brown J, Li S, Swanson J, Rayner JC, Jiang RHY, Adams JH. 2018. Uncovering the essential genes of the human malaria parasite Plasmodium falciparum by saturation mutagenesis. Science 360.

35. Cilingir G, Broschat SL, Lau AO. 2012. ApicoAP: the first computational model for identifying apicoplast-targeted proteins in multiple species of Apicomplexa. PLoS One 7:e36598.

36. Abrahamsen MS, Templeton TJ, Enomoto S, Abrahante JE, Zhu G, Lancto CA, Deng M, Liu C, Widmer G, Tzipori S, Buck GA, Xu P, Bankier AT, Dear PH, Konfortov BA, Spriggs HF, Iyer L, Anantharaman V, Aravind L, Kapur V. 2004. Complete genome sequence of the apicomplexan, Cryptosporidium parvum. Science 304:441–5.

37. Xu P, Widmer G, Wang Y, Ozaki LS, Alves JM, Serrano MG, Puiu D, Manque P, Akiyoshi D, Mackey AJ, Pearson WR, Dear PH, Bankier AT, Peterson DL, Abrahamsen MS, Kapur V, Tzipori S, Buck GA. 2004. The genome of Cryptosporidium hominis. Nature 431:1107–12.

38. Zhu G, Marchewka MJ, Keithly JS. 2000. Cryptosporidium parvum appears to lack a plastid genome. Microbiology (Reading) 146 (Pt 2):315–321.

39. Bylund GO, Wipemo LC, Lundberg LA, Wikstrom PM. 1998. RimM and RbfA are essential for efficient processing of 16S rRNA in Escherichia coli. J Bacteriol 180:73–82.

40. Liu K, Lee KP, Duan J, Kim EY, Singh RM, Di M, Meng Z, Kim C. 2023. Cooperative role of AtRsmD and AtRimM proteins in modification and maturation of 16S rRNA in plastids. Plant J 114:310–324.

41. van Dooren GG, Striepen B. 2013. The algal past and parasite present of the apicoplast. Annu Rev Microbiol 67:271–89.

42. Leveque MF, Berry L, Cipriano MJ, Nguyen HM, Striepen B, Besteiro S. 2015. Autophagy-Related Protein ATG8 Has a Noncanonical Function for Apicoplast Inheritance in Toxoplasma gondii. mBio 6:e01446–15.

43. Niu Z, Ye S, Liu J, Lyu M, Xue L, Li M, Lyu C, Zhao J, Shen B. 2022. Two apicoplast dwelling glycolytic enzymes provide key substrates for metabolic pathways in the apicoplast and are critical for Toxoplasma growth. PLoS Pathog 18:e1011009.

44. Sanchez SG, Bassot E, Cerutti A, Mai Nguyen H, Aida A, Blanchard N, Besteiro S. 2023. The apicoplast is important for the viability and persistence of Toxoplasma gondii bradyzoites. Proc Natl Acad Sci U S A 120:e2309043120.

45. McFadden GI, Yeh E. 2017. The apicoplast: now you see it, now you don’t. Int J Parasitol 47:137–144.

46. Huynh MH, Carruthers VB. 2009. Tagging of endogenous genes in a Toxoplasma gondii strain lacking Ku80. Eukaryot Cell 8:530–9.

47. Sidik SM, Huet D, Lourido S. 2018. CRISPR-Cas9-based genome-wide screening of Toxoplasma gondii. Nat Protoc 13:307–323.

48. Shen B, Brown KM, Lee TD, Sibley LD. 2014. Efficient gene disruption in diverse strains of Toxoplasma gondii using CRISPR/CAS9. mBio 5:e01114–14.

49. Alvarez-Jarreta J, Amos B, Aurrecoechea C, Bah S, Barba M, Barreto A, Basenko EY, Belnap R, Blevins A, Bohme U, Brestelli J, Brown S, Callan D, Campbell LI, Christophides GK, Crouch K, Davison HR, DeBarry JD, Demko R, Doherty R, Duan Y, Dundore W, Dyer S, Falke D, Fischer S, Gajria B, Galdi D, Giraldo-Calderon GI, Harb OS, Harper E, Helb D, Howington C, Hu S, Humphrey J, Iodice J, Jones A, Judkins J, Kelly SA, Kissinger JC, Kittur N, Kwon DK, Lamoureux K, Li W, Lodha D, MacCallum RM, Maslen G, McDowell MA, Myers J, Nural MV, Roos DS, et al. 2024. VEuPathDB: the eukaryotic pathogen, vector and host bioinformatics resource center in 2023. Nucleic Acids Res 52:D808–D816.

50. Sievers F, Wilm A, Dineen D, Gibson TJ, Karplus K, Li W, Lopez R, McWilliam H, Remmert M, Soding J, Thompson JD, Higgins DG. 2011. Fast, scalable generation of high-quality protein multiple sequence alignments using Clustal Omega. Mol Syst Biol 7:539.

51. Yu G. 2020. Using ggtree to Visualize Data on Tree-Like Structures. Curr Protoc Bioinformatics 69:e96.

52. Plattner F, Yarovinsky F, Romero S, Didry D, Carlier MF, Sher A, Soldati-Favre D. 2008. Toxoplasma profilin is essential for host cell invasion and TLR11-dependent induction of an interleukin-12 response. Cell Host Microbe 3:77–87.

