## Supplemental Figure for "*In vivo* CRISPR screens identify novel virulence genes among proteins of unassigned subcellular localization in *Toxoplasma*"

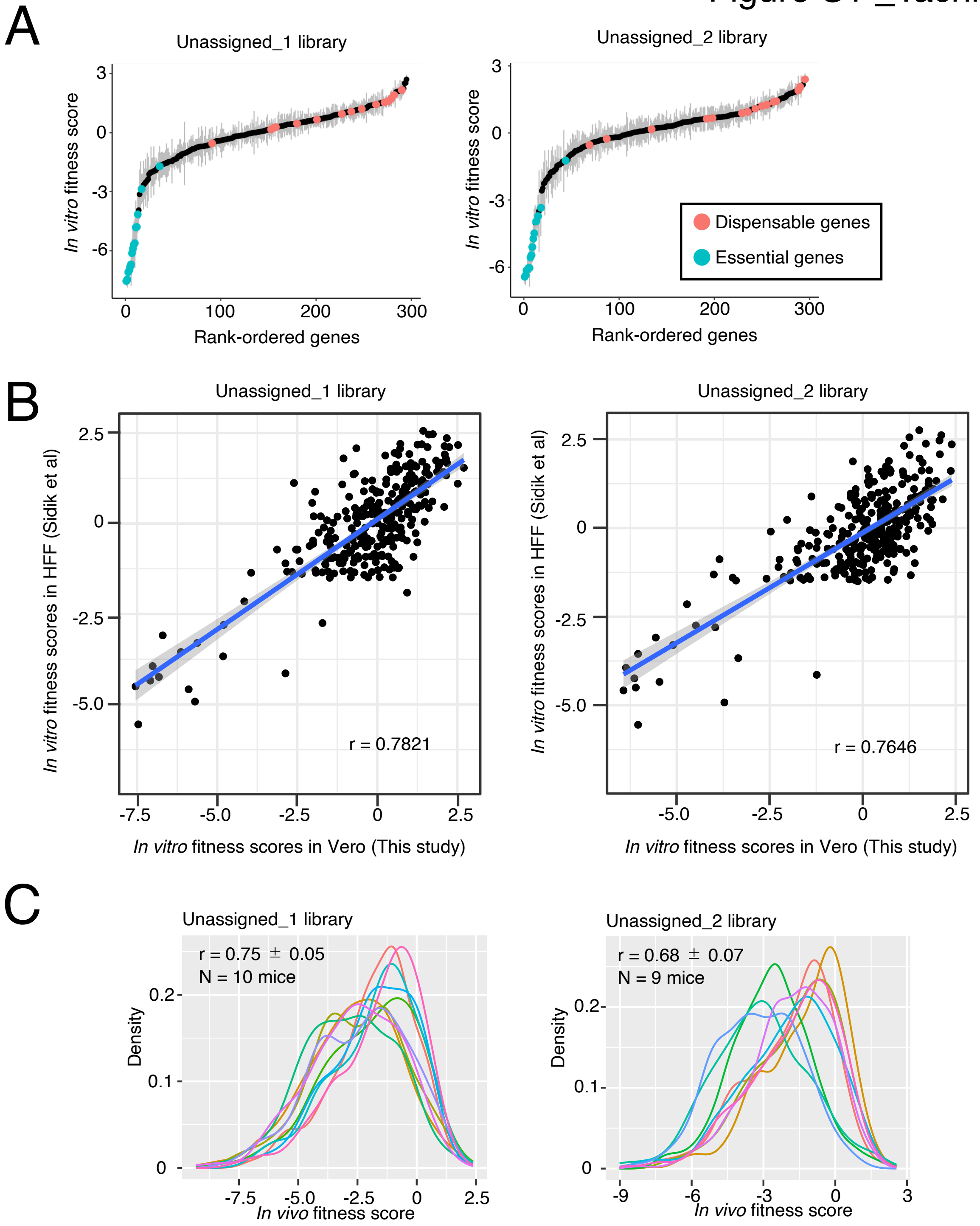


**Supplementary Figure 1. Assessing the reproducibility of CRISPR screens**

**Related to Figure 1.**

(A) Rank-ordered plots for *in vitro* fitness scores of 4$^{th}$ passage. (B) Correlation between our *in vitro* fitness scores and the *in vitro* fitness scores in HFF. (C) Overlay of *in vivo* fitness scores for each mouse. Pearson's correlation coefficients are shown as mean ± SD.

Figure S2 _Tachibana et al.

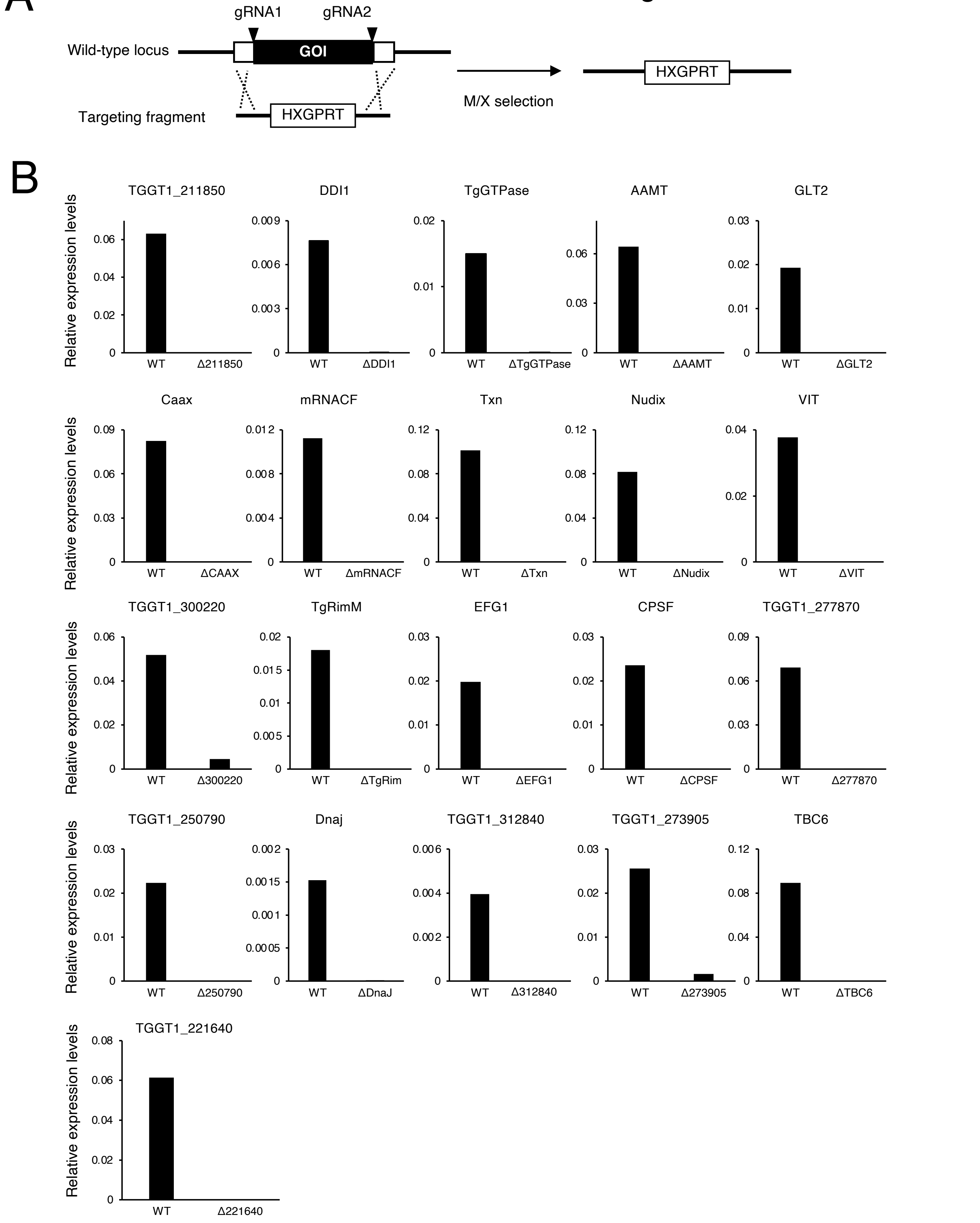


**Supplementary Figure 2. Generating gene knockout parasites. Related to Figure 2.**

(A) Schematic of gene knockout strategy. (B) Quantitative RT-PCR validations for indicated parasites.

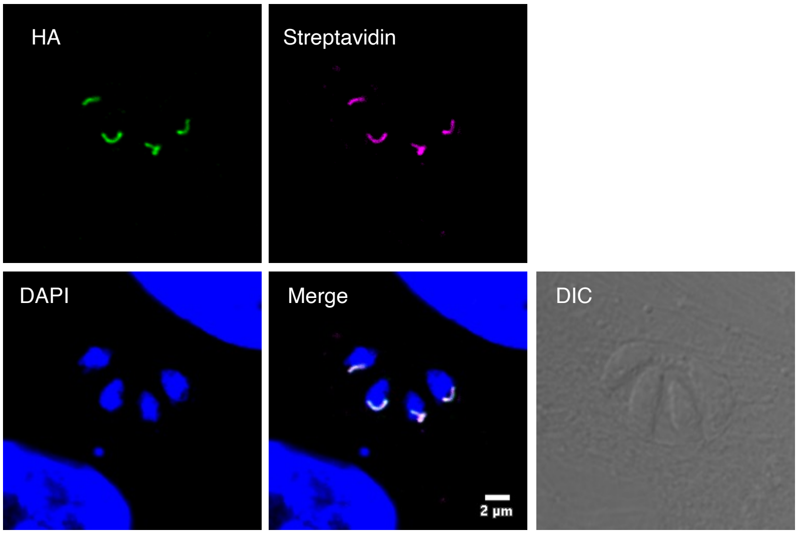


**Supplementary Figure 3. TgRimM$^{\Delta ZF}$-HA localized in the apicoplast.**

**Related to Figure 4.**

Immunofluorescence images of ΔTgRimM::TgRimM$^{\Delta ZF}$-HA showing TgRimM$^{\Delta ZF}$-HA (green) is co-stained with streptavidin (magenta), an apicoplast marker. Data are representative of three independent experiments.
